# Defining the genetic control of human blood plasma N-glycome using genome-wide association study

**DOI:** 10.1101/365486

**Authors:** Sodbo Zh. Sharapov, Yakov A. Tsepilov, Lucija Klaric, Massimo Mangino, Gaurav Thareja, Mirna Simurina, Concetta Dagostino, Julia Dmitrieva, Marija Vilaj, Frano Vuckovic, Tamara Pavic, Jerko Stambuk, Irena Trbojevic-Akmacic, Jasminka Kristic, Jelena Simunovic, Ana Momcilovic, Harry Campbell, Malcolm Dunlop, Susan Farrington, Maja Pucic-Bakovic, Christian Gieger, Massimo Allegri, Edouard Louis, Michel Georges, Karsten Suhre, Tim Spector, Frances MK Williams, Gordan Lauc, Yurii Aulchenko

## Abstract

Glycosylation is a common post-translational modification of proteins. It is known, that glycans are directly involved in the pathophysiology of every major disease. Defining genetic factors altering glycosylation may provide a basis for novel approaches to diagnostic and pharmaceutical applications. Here, we report a genome-wide association study of the human blood plasma N-glycome composition in up to 3811 people. We discovered and replicated twelve loci. This allowed us to demonstrate a clear overlap in genetic control between total plasma and IgG glycosylation. Majority of loci contained genes that encode enzymes directly involved in glycosylation (*FUT3/FUT6, FUT8, B3GAT1, ST6GAL1, B4GALT1, ST3GAL4, MGAT3*, and *MGAT5*). We, however, also found loci that are likely to reflect other, more complex, aspects of plasma glycosylation process. Functional genomic annotation suggested the role of *DERL3*, which potentially highlights the role of glycoprotein degradation pathway, and such transcription factor as *IKZF1*.

## Introduction

Glycosylation - addition of carbohydrates to a substrate - is a common cotranslational and posttranslational modification of proteins. that affects the physical properties of proteins (solubility, conformation, folding, stability, trafficking, etc.) [1–4] as well as their biological functions - from protein-protein interactions, interaction of proteins with receptors, to cell-cell, cell-matrix, and host-pathogen interactions [2,3,5,6]. It has been estimated that more than half of all proteins are glycosylated [7–9]. Given the fact that glycans participate in many biological processes, it is therefore not surprising that molecular defects in protein glycosylation pathways are increasingly recognized as direct causes of diseases, such as rheumatoid arthritis, cardiometabolic disorders, cancer, variety of autoimmune diseases, type 2 diabetes, inflammatory bowel disease and others [10–17]. More specifically, a variety of N-glycan structures are now considered as disease markers and represent diagnostic as well as therapeutic targets [5,12,18–25]. Defining the genetic control of protein glycosylation expands our knowledge about the regulation of this fundamental biological process, and it may also shed new light onto how alterations in glycosylation can lead to the development of complex human diseases [11].

Previous genome-wide association studies (GWAS) of total plasma protein N-glycome measured with high performance liquid chromatography (HPLC) discovered six loci associated with protein glycosylation [26,27]. Four of these contained genes that have well characterized roles in glycosylation: the fucosyltransferases *FUT6* and *FUT8*, glucuronyltransferase *B3GAT1*, and glucosaminyltransferase *MGAT5*. Other two loci—one near *SLC9A9* on chromosomes 3 and one near *HNF1a* on chromosome 12—did not contain any genes known to be involved in glycosylation processes. A functional *in vitro* follow-up study in HepG2 cells [27] on the *HNF1a* locus on chromosome 12, showed that its gene product acts as a co-regulator of expression of most fucosyltransferase genes (*FUT3, FUT5, FUT6, FUT8, FUT10, FUT11*). In addition, it co-regulates expression of genes encoding key enzymes required for the synthesis of GDP-fucose, the substrate of these fucosyltransferases. It was concluded that *HNF1a* is one of the master regulators of protein glycosylation, influencing both core and antennary fucosylation [26]. The locus on chromosome 3 contained *SLC9A9*, a gene that encodes a proton pump which affects pH in the endosomal compartment, reminiscent of recent findings that changes in Golgi pH can impair protein sialylation, suggesting a possible mechanism for the observed association with N-glycosylation traits.

Since 2011, when the latest GWAS of plasma N-glycome was published, new technologies for glycome profiling were developed [28]. Ultra-performance liquid chromatography (UPLC) became a widely used technology for accurate analysis of plasma N-glycosylation due to its superior sensitivity, resolution, speed, and its capability to provide branch-specific information of glycan structures [29]. Moreover, new imputation panels (such as 1000 Genomes [30] and HRC [31]) became available, increasing the resolution and power of genetic mapping.

In this work, we aimed to advance our understanding of the genetic control of the human plasma N-glycome, and to establish a public resource that will facilitate future studies linking glycosylation and complex human diseases. For that, we performed and reported results of GWAS on 113 plasma glycome traits measured by UPLC and genotypes imputed to the 1000 Genomes reference panel in 2,763 participants of TwinsUK. Further we replicated our findings in 1,048 samples from three independent and genetically diverse cohorts - PainOR, SOCCS and QMDiab.

## Results

### Replication of previously reported loci

We started with replication of six loci that were reported previously. Huffman and colleagues [27] analyzed four independent cohorts with total sample size of 3,533, using plasma N-glycome measured with HPLC. Because of technological differences, there is no one-to-one correspondence between HPLC and UPLC traits, and exact replication is not possible. Therefore, we analyzed association of the SNPs reported by [27] with all 113 UPLC traits measured in this study, and considered a locus replicated if we observed P-value ≤ 0.05/(6×30) = 2.78 × 10^−4^ (where 30 is a number of principal components, explaining 99% of the variation of the 113 studied traits) in the TwinsUK cohort (N=2,763). Using this procedure, we replicated 5 of the 6 previously reported SNPs (Table 1). For more details, see Supplementary Table 1.

**Table 1.** Replication of six previously found loci (Huffman et al, 2011). Replicated loci are in bold. CHR:POS - chromosome and position of SNP, Eff/Ref - effective and reference allele, Gene - candidate gene for the locus, EAF - effective allele frequency, N - sample size, BETA(SE) - effect and standard error of effect estimates, min P – minimal P-value, observed across glycomic traits.

| SNP | CHR:POS | Gene | Eff/Ref | Results of Huffman et al, 2011 (N=3,533) |  |  |  | This study, TwinsUK (N=2,763) |  |  |
| --- | --- | --- | --- | --- | --- | --- | --- | --- | --- | --- |
|  |  |  |  | EAF | N | BETA(SE) | min P | EAF | BETA(SE) | min P |
| **rs1257220** | 2:135015347 | *MGAT5* | A/G | 0.26 | 3263 | 0.19(0.03) | 1.80E-10 | 0.25 | 0.16(0.032) | 3.98E-07 |
| **rs4839604** | 3:142960273 | *SLC9A9* | C/T | 0.77 | 3320 | -0.22(0.03) | 3.50E-13 | 0.83 | -0.11(0.038) | 2.50E-03 |
| **rs7928758** | 11:134265967 | *B3GAT1* | T/G | 0.88 | 3233 | 0.23(0.04) | 1.66E-08 | 0.84 | 0.24(0.038) | 6.07E-10 |
| **rs735396** | 12:121438844 | *HNF1a* | T/C | 0.61 | 3236 | 0.18(0.03) | 7.81E-12 | 0.65 | 0.18(0.031) | 6.06E-09 |
| **rs11621121** | 14:65822493 | *FUT8* | C/T | 0.43 | 3234 | 0.27(0.03) | 1.69E-23 | 0.40 | 0.21(0.029) | 1.60E-12 |
| **rs3760776** | 19:5839746 | *FUT6* | G/A | 0.87 | 3262 | 0.44(0.04) | 3.18E-29 | 0.91 | 0.56(0.050) | 3.71E-28 |

These results not only confirm previous and establish five plasma glycome loci as replicated, but also demonstrate that our study is well powered (among replicated loci, all P-value were less than 4 × 10^−7^).

### Discovery and replication of new loci

The discovery cohort comprised 2,763 participants of the TwinsUK study with genotypes available for 8,557,543 SNPs. The genomic control inflation factor varied from 0.99 to 1.02, suggesting that influences of residual population stratification on the test statistics were small (see Supplementary Table 2; QQ-plots in Supplementary Figure 1). In total, 906 SNPs located in 14 loci were significantly associated (P-value ≤ 5 × 10^−8^ / 30 = 1.66 × 10^−9^, where 30 is a number of principal components, explaining 99% of the variation of the 113 studied traits) with at least one of 113 glycan traits (in total 5,052 SNP-trait associations, see Figure 1, Table 1). Out of 113 traits, 68 were significantly associated with at least one of the 14 loci. For more details, see Supplementary Table 3.

**Figure 1.**
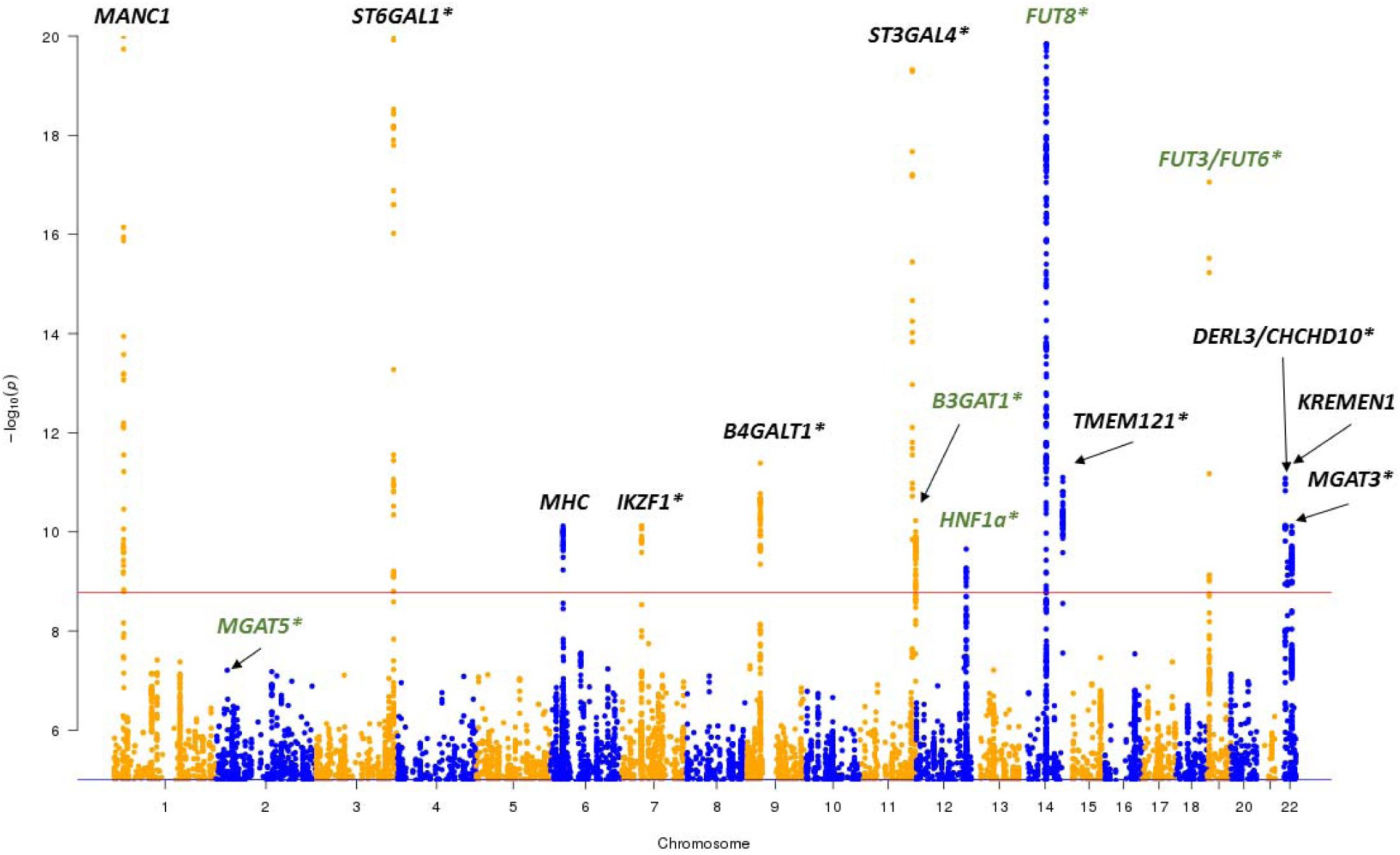
Manhattan plot of discovery GWAS (after correction for genomic control). Red line corresponds to the genome-wide significance threshold of 1.7 × 10^−9^. For each SNP the lowest P-value among 113 traits is shown. Only SNPs with P-values ≤ 1 × 10^−5^ are shown. Points with −log10(P-value)>20 are depicted at −log10 (P-value)=20. Green colored gene labels marks loci that were found in previous GWAS [27]; black colored marks novel loci, * - replicated loci.

Among fourteen loci, four were previously reported as associated with the plasma N-glycome. Three loci—on chromosome 12 at 121 Mb (leading SNP rs1169303, intronic variant of the *HNF1a* gene), on chromosome 14 at 105 Mb (leading SNP - rs7147636 located in the intron of *FUT8* gene), and on chromosome 19 at 58 Mb (leading SNP: rs7255720, upstream variant of the *FUT6* gene)—were reported to be associated with the plasma N-glycome in two previous GWAS [26,27], while association of a locus on chromosome 11 at 126 Mb (leading SNP - rs1866767 located in the intron of *B3GAT1* gene) was reported only in the latest GWAS meta-analysis of plasma N-glycome [27].

Ten further loci that have not been reported before were found here. In order to replicate our findings, we have performed association analysis of these ten SNPs in three independent cohorts— PainOR, SOCCS and QMDiab (total N =1,048)—and then meta-analyzed the results. Seven of ten novel loci were replicated at threshold P-value ≤ 0.05/10 = 0.005 (see Table 2). The direction of association was concordant between discovery and replication for all ten loci. The effects of loci between the replication cohorts were homogeneous (P-value of Cochran’s Q-test varied from 0.07 to 0.96, see Supplementary Table 3).

**Table 2.** Fourteen loci genome-wide significantly associated with at least one of the 113 traits in this study. Ten loci in the upper part of the table are novel, and four loci in the lower part of the table were found previously. Replicated loci are in bold. CHR:POS - chromosome and position of SNP, Gene –suggested candidate genes (see Table 3), Eff/Ref - effective and reference allele, EAF - effective allele frequency, BETA(SE) - effect and standard error of effect estimates, P-value - P-value after GC correction, Top trait –glycan trait with the strongest association (lowest P-value), N traits - total number of traits significantly associated with given locus, N - sample size of replication.

| SNP | CHR:POS | Gene | Eff / Ref | Discovery |  |  |  |  | Replication |  |  |  |
| --- | --- | --- | --- | --- | --- | --- | --- | --- | --- | --- | --- | --- |
|  |  |  |  | EAF | BETA (SE) | P-value | Top trait | N traits | EAF | BETA (SE) | P | N |
| Novel loci |  |  |  |  |  |  |  |  |  |  |  |  |
| rs186127900 | 1:25318225 | - | G/T | 0.99 | -1.26 (0.119) | 4.04E-24 | PGP82 | 26 | 0.99 | -0.35 (0.224) | 1.22E-01 | 1093 |
| rs59111563 | 3:186722848 | ST6GAL1 | D/I | 0.74 | 0.34 (0.031) | 1.09E-26 | PGP41 | 3 | 0.73 | 0.32 (0.048) | 9.50E-12 | 1088 |
| rs3115663 | 6:31601843 | - | T/C | 0.80 | 0.26 (0.040) | 7.65E-11 | PGP18 | 1 | 0.83 | 0.06 (0.059) | 3.01E-01 | 1093 |
| rs6421315 | 7:50355207 | IKZF1 | G/C | 0.59 | 0.19 (0.029) | 7.57E-11 | PGP60 | 2 | 0.60 | 0.27 (0.043) | 5.67E-10 | 1077 |
| rs13297246 | 9:33128617 | B4GALT1 | G/A | 0.83 | -0.26 (0.038) | 4.11E-12 | PGP67 | 2 | 0.83 | -0.26 (0.059) | 8.66E-06 | 1093 |
| rs3967200 | 11:126232385 | ST3GAL4 | C/T | 0.88 | -0.49 (0.043) | 1.51E-27 | PGP17 | 7 | 0.86 | -0.53 (0.062) | 6.85E-18 | 1093 |
| rs35590487 | 14:105989599 | IGH /<br>TMEM121 | C/T | 0.77 | -0.24 (0.034) | 7.98E-12 | PGP62 | 2 | 0.78 | -0.17 (0.058) | 3.67E-03 | 1093 |
| rs9624334 | 22:24166256 | DERL3 /<br>CHCHD10 | G/C | 0.85 | 0.28 (0.040) | 8.38E-12 | PGP63 | 2 | 0.86 | 0.42 (0.062) | 2.09E-11 | 1086 |
| rs140053014 | 22:29550678 | - | I/D | 0.98 | -0.67 (0.106) | 4.05E-10 | PGP109 | 1 | 0.98 | -0.24 (0.165) | 1.50E-01 | 1079 |
| rs909674 | 22:39859169 | MGAT3 | C/A | 0.27 | 0.22 (0.033) | 7.72E-11 | PGP56 | 3 | 0.25 | 0.18 (0.053) | 5.70E-04 | 1045 |
| Previously implicated loci |  |  |  |  |  |  |  |  |  |  |  |  |
| rs1866767 | 11:134274763 | B3GAT1 | C/T | 0.87 | 0.28 (0.043) | 5.95E-11 | PGP33 | 3 |  |  |  |  |
| rs1169303 | 12:121436376 | HNF1a | A/C | 0.51 | 0.19 (0.029) | 2.23E-10 | PGP30 | 2 |  |  |  |  |
| rs7147636 | 14:66011184 | FUT8 | T/C | 0.33 | -0.39 (0.030) | 6.63E-37 | PGP20 | 17 |  |  |  |  |
| rs7255720 | 19:5828064 | FUT6 | G/C | 0.96 | 1.14 (0.068) | 2.53E-55 | PGP110 | 18 |  |  |  |  |

Given seven replicated novel loci found in this study and five loci found previously and replicated in this study we now have 12 replicated loci in total.

## Functional annotation *in-silico*

### Analysis of possible effects of genetic variants with VEP

We have used variant effect predictor [32] in order to find functional variants that potentially disturb amino acid sequence and may explain association in some loci. For that, within each locus, we identified a set of SNPs that are likely to contain the functional variant by selecting SNPs that had association p-value deviating from the minimal p-value by less than one order of magnitude. The results of variant effect predictor [32] annotation of the resulting 214 SNPs are presented in Supplementary Table 4b. For the locus on chromosome 19 at 58 Mb, we have observed that rs17855739 variant is missense for five transcripts of the *FUT6* gene; for four of these transcripts rs17855739 was classified as probably damaging and for one as benign by PolyPhen [33], whereas SIFT [34] classified all five variants as deleterious. For the locus on chromosome 22 at 24 Mb, we detected rs3177243 variant that is missense for three transcripts of *DERL3* gene, for which rs3177243 was predicted to be deleterious by SIFT (although was classified as benign by PolyPhen). For locus on chromosome 14 at 105/106 Mb, in *TMEM121* gene, we observed that rs10569304 variant led to three nucleotides in-frame deletion in two transcripts of *TMEM121* gene.

### Gene-set and tissue/cell enrichment analysis

For prioritizing genes in associated regions (based on their predicted function) and gene set and tissue/cell type enrichment analyses we used DEPICT software [35]. When running DEPICT analyses on the 14 genome-wide significant loci (from Table 1) we identified tissue/cell type enrichment (with FDR<0.05) for six tissue/cell types: plasma cells, plasma, parotid gland, salivary glands, antibody producing cells and B-lymphocytes (see Supplementary Table 5c). We did not identify any significant enrichment for gene-sets (all FDR > 0.2, Supplementary Table 5b). Based on predicted gene function and reconstituted gene sets, DEPICT suggestively prioritized three genes - *FUT3, DERL3* and *FUT8* for three loci (on chromosome 19 at 58 Mb, on chromosome 22 at 24 Mb and on chromosome 14 at 65/66 Mb) with FDR < 0.20 (see Supplementary Table 5a). We have also analyzed 93 loci with P-value ≤ 1 × 10^−5^/30 (Supplementary Table 6), however, all results had FDR > 0.2.

### Overlap with complex traits

We next investigated the potential pleiotropic effects of our loci on other complex human traits and diseases, using PhenoScanner v1.1 database [36]. For twelve replicated SNPs (Table 1 and Table 2), we looked up traits that were genome-wide significantly (P-value≤ 5 × 10^−8^) associated with the same SNP or a SNP in strong (r^2^< 0.7) linkage disequilibrium. The results are summarized in Supplementary Table 7. For eight out of twelve loci, we observed associations with a number of complex traits. Four loci (near *IKZF1, FUT8, MGAT3* and *DERL3*) were associated with levels of glycosylation of immunoglobulin G (IgG) [13]. Two loci (on chromosome 12 at 121 Mb and on chromosome 11 at 126 Mb, containing *HNF1a* and *ST3GAL4* genes respectively) were associated with LDL and total cholesterol levels [37,38]. The locus containing *HNF1a* was additionally associated with level of plasma C reactive protein [39,40] and gamma glutamyl transferase level [41]. Locus on chromosome 22 at 39 Mb (containing *MGAT3*) was associated with adult height [42]. Locus on chromosome 14 at 65/66 Mb (near *FUT8*) was associated with age at menarche [43]. Note, however, that PhenoScanner analysis does not allow distinguishing between pleiotropy of a variant shared between traits, and linkage disequilibrium between different functional variants affecting separate traits.

### Pleiotropy with eQTLs

We next attempted to identify genes whose expression levels could potentially mediate the association between SNPs and plasma N-glycome. For this we performed a summary-data based Mendelian randomization (SMR) analysis followed by heterogeneity in dependent instruments (HEIDI) analysis [44] using a collection of eQTL data from a range of tissues, including blood [45], 44 tissues as provided in the GTEx database version 6p [46] and six blood cell lines collected in the CEDAR study (see Supplementary Note 3 and [47]) - five immune cell populations (CD4+, CD8+, CD19+, CD14+, CD15+) and platelets. In short, SMR test aims at testing the association between gene expression (in a particular tissue) and a trait using the top associated SNP as a genetic instrument. Significant SMR test may indicate that the same functional variant determines both expression and the trait of interest (causality or pleiotropy), but may also indicate the possibility that functional variants underlying gene expression are in linkage disequilibrium with those controlling the traits. Inferences whether functional variant may be shared between plasma glycan trait and expression were made based on HEIDI test: P_HEIDI_≥ 0.05 (likely shared), 0.05 >P_HEIDI_≥ 0.001 (possibly shared), P_HEIDI_< 0.001 (sharing is unlikely).

We applied SMR/HEIDI analyses for replicated loci that demonstrated genome-significant association in our discovery data (11 loci in total). In total, we included in the analysis expression levels of 20,448 transcripts (probes). For fifteen probes located in seven loci associated with plasma glycosylation we observed significant (P_SMR_≤ 0.05/20,448=2.445 × 10^−6^) association to the top SNPs associated with plasma N-glycome (see Supplementary Table 8). Subsequent HEIDI test showed that the hypothesis of shared functional variant between plasma glycan traits and expression is most likely (P_HEIDI_>0.05) for four probes: *ST6GAL1* in whole blood (from Westra *et al*., [45]; *TMEM121* in whole blood (GTEx, [46]); *MGAT3* in CD19+ cell line (CEDAR, [47]) and *CHCHD10* in whole blood (results of Westra *et al*., [45]). For other five probes we conclude that the functional variant is possibly shared (0.001<P_HEIDI_<0.05) between glycan traits and expression of *ST3GAL4* (in two different tissues: muscle skeletal and pancreas, GTEx [46]); *B3GAT1* (in two tissues: whole blood from Westra *et al*., [45], and lung tissue from GTEx, [46]); *SYNGR1* (in tibial nerve tissue from GTEx [46]).

### Summary of in-silico follow-up

We compared the genes suggested by our *in silico* functional investigation with candidate genes suggested previously for five known loci (see Table 3). For three out of five loci (*B3GAT1, FUT8, FUT6/FUT3*) we selected the same genes as suggested by the authors of previous study [27]. All three genes are known to be involved into the glycan synthesis pathways. The *FUT8* locus was associated mostly with core-fucosylated biantennary glycans, that are known to be linked to the immunoglobulins [9]. As *FUT8* gene codes fucosyltransferase 8, an enzyme responsible for the addition of core fucose to glycans, this gene is most biologically plausible in this locus. *FUT3* and *FUT6* encode fucosyltransferases 3 and 6 that catalyze the transfer of fucose from GDP-beta-fucose to alpha-2,3 sialylated substrates. The *FUT3/FUT6* locus was associated with antennary fucosylation of tri- and tetra-antennary sialylated glycans, and therefore we consider these genes as good candidates. Moreover, in the *FUT6* gene (chromosome 19, 58 Mb) we found missense variant rs17855739 (substitution G>A) that leads to amino acid change from negatively charged glutamic acid to positively charged lysine. PolyPhen and SIFT predicted this variant as deleterious for transcripts of *FUT6* gene. Thus, we can consider this SNP as possible causal functional variant.

**Table 3.** Summary of functional *in-silico* annotation for the replicated loci. For each locus we report the gene nearest to the top SNP and a plausible candidate gene with sources of evidence. CV - coding variant for the gene, suggested by VEP; SMR/HEIDI – evidence by pleiotropy with expression by SMR-HEIDI; D – evidence by DEPICT; Funct. studies – evidence by functional studies; Glycan synth. –known glycan synthesis gene in the locus; Prev. annot. – the region was previously implicated in the glycome GWAS, and the gene was suggested as candidate (P1 - gene was reported as affecting plasma N-glycome by Huffman et al., 2011 [27]; IgG - gene was reported as affecting IgG glycome either by Lauc et al., 2013 [13] and/or by Shen et al., 2017 [48]).

| Locus | Nearest gene | Candidate Gene | CV | SMR/HEIDI | DEPICT | Func. studies | Glycan synth. | Prev. annot. |
| --- | --- | --- | --- | --- | --- | --- | --- | --- |
| **Previously implicated loci** |  |  |  |  |  |  |  |  |
| 2:135015347 |  | *MGAT5* | - |  |  |  | + | PI |
| 11:134274763 | *B3GAT1* | *B3GAT1* | - | whole blood/<br>lung |  |  | + | PI |
| 12:121436376 | *HNF1a* | *HNF1a* | - |  |  | [27] |  | PI |
| 14:66011184 | *FUT8* | *FUT8* | - |  | FDR<20% |  | + | PI,<br>IgG |
| 19:5828064 | *NRTN* | *FUT3* | - |  | FDR<20% |  | + | PI |
|  | *NRTN* | *FUT6* | rs17855739 |  |  |  | + | PI,<br>IgG |
| **Novel loci** |  |  |  |  |  |  |  |  |
| 3:186722848 | *ST6GAL1* | *ST6GAL1* | - | whole blood |  |  | + | IgG |
| 7:50355207 | *IKZF1* | *IKZF1* | - |  |  |  |  | IgG |
| 9:33128617 | *B4GALT1* | *B4GALT1* | - |  |  |  | + | IgG |
| 11:126232385 | *ST3GAL4* | *ST3GAL4* | - | muscle<br>skeletal/pancreas |  |  | + |  |
| 14:105989599 | *C14orf80* | *TMEM121* | rs10569304 | whole blood |  |  |  |  |
|  | *C14orf80* | *IGH* |  |  |  |  |  | IgG |
| 22:24166256 | *SMARCB1* | *DERL3* | rs3177243 |  | FDR<20% |  |  | IgG |
|  | *SMARCB1* | *CHCHD10* | - | whole blood |  |  |  |  |
| 22:39859169 | *MGAT3* | *MGAT3* | - | CD19+ (B cells) |  |  | + | IgG |

For the other two loci (on chromosome 3 at 142 Mb and on chromosome 12 at 121 Mb) we were not able to prioritize genes by VEP, DEPICT, and eQTL analyses. However, the first locus contained *MGAT5* gene coding mannosyl-glycoprotein-N-acetyl glucosaminyl-transferase that is involved into the glycan synthesis pathways. The second locus contained several genes including *HNF1a* which was previously shown to co-regulate the expression of most fucosyltransferase (*FUT3, FUT5, FUT6, FUT8, FUT10, FUT11*) genes in a human liver cancer cell line (HepG2 cells); as well as to co-regulating gene expression levels of key enzymes needed for synthesis of GDP-fucose, the substrate for fucosyltransferases, thereby regulating multiple stages in the fucosylation process [26]. Thus, we considered *HNF1a* as the candidate gene for this locus.

Four of the seven novel loci contain genes that are known to be involved in glycan synthesis pathways - *ST6GAL1, ST3GAL4, B4GALT1* and *MGAT3* (see Table 3). Moreover, summary level Mendelian randomization (SMR) and HEIDI analyses have shown that expression of *ST6GAL1* and *MGAT3* genes may mediate the association between corresponding loci and plasma N-glycome. *ST6GAL1* and *ST3GAL4* genes encode sialyltransferases, enzymes which catalyze the addition of sialic acid to various glycoproteins. The locus containing *ST6GAL1* was associated with ratio of sialylated and non-sialylated galactosylated biantennary glycans. The locus containing *ST3GAL4* was associated with galactosylated sialylated tri- and tetra-antennary glycans. The locus containing *MGAT3* was associated with proportion of bisected biantennary glycans. This latter gene encodes the enzyme N-acetylglucosaminyltransferase, which is responsible for the addition of bisecting GlcNAc. The *B4GALT1* gene encodes galactosyltransferase, which adds galactose during the biosynthesis of different glycoconjugates. This gene was associated with galactosylation of biantennary glycans. Thus, we observe consistency between known enzymatic activities of the products of selected candidate genes and the spectrum of glycans that are associated with corresponding loci.

The other three novel loci do not contain genes that are known to be directly involved in glycan synthesis. Variant rs9624334 (chromosome 22 at 24 Mb) is located in the intron of *SMARCB1* gene that is known to be important in antiviral activity, inhibition of tumor formation, neurodevelopment, cell proliferation and differentiation [49]. However, gene prioritization analysis (DEPICT) showed, that the possible candidate gene is *DERL3*, which encodes a functional component of endoplasmic reticulum (ER)-associated degradation for misfolded luminal glycoproteins [50] (see Table 3). Additionally, VEP analysis demonstrated that the leading rs9624334 variant in this locus is in strong LD (R^2^=0.98 in 1000 Genome EUR samples) with rs3177243, which is a *DERL3* coding variant predicted to be deleterious by SIFT and PolyPhen. However, the SMR/HEIDI analysis suggested that the association with N-glycome could be (also) mediated by expression of *CHCHD10* gene, which encodes a mitochondrial protein that is enriched at cristae junctions in the intermembrane space. The *CHCHD10* gene has the highest expression in heart and liver and the lowest expression in spleen [51]. While the role of mitochondrial proteins in glycosylation processes remains speculative, we propose *CHCHD10* as a candidate based on our eQTL pleiotropy analysis. Thus, we consider two genes - *DERL3* and *CHCHD10* - as possible candidate genes at this locus. Interestingly, this and the *MGAT3* loci were associated with similar glycan traits (core-fucosylation of bisected glycans). This indicates that core fucosylation of bisected glycans is under joint control of *MGAT3* and *DERL3/CHCHD10*.

The locus on chromosome 14 at 105 Mb contains the *IGH* gene that encodes immunoglobulin heavy chains. This locus is associated with sialylation of core-fucosylated biantennary monogalactosylated structures that are biochemically close to those affected by *ST6GAL1* gene. As IgG is the most prevalent glycosylated plasma protein [9], one would consider *IGH* as a good candidate, as indeed was suggested by Shen and colleagues [48]. However, our functional annotation results (SMR/HEIDI and VEP) suggest that association of this locus with plasma N-glycome may be mediated by regulation of expression of *TMEM121* gene. This gene encodes transmembrane protein 121 that is highly expressed in heart as well as being detected in pancreas, liver and skeletal muscle. Moreover, for the lead SNP rs35590487 we found a variant rs10569304 that is in strong linkage disequilibrium (R^2^ =0.95 In 1000 Genome EUR samples) with it, and which leads to inframe deletion in protein coding region of the *TMEM121* gene. Therefore, we consider two genes *–IGH* and *TMEM121–* as candidate genes for this locus.

For the locus on chromosome 7 at 50 Mb we were not able to select a candidate gene based on results of our *in-silico* functional annotation. This locus was previously reported to be associated with glycan levels of IgG [13], and authors suggested that *IKZF1* may be considered as a candidate gene in the region. The *IKZF1* gene codes the DNA-binding protein Ikaros that acts as a transcriptional regulator and is associated with chromatin remodeling. It is considered an important regulator of lymphocyte differentiation. Taking into account that IgG (the most abundant glycoprotein in the blood plasma [9]) are secreted by B cells [52], *IKZF1* seems to be a plausible candidate gene.

To identify possible clusters in the gene network of plasma N-glycosylation we draw a graph in which eleven genome-wide significant loci and genome-wide significantly (P-value ≤ 1.66 × 10^−9^) associated glycan traits were presented as nodes, and edges represent observed significant associations (see Figure 2). We labeled each glycan trait as “immunoglobulin-linked” (Ig-linked), “non-immunoglobulin-linked” (non-Ig-linked) or mixed (could be linked to either) depending on the contribution of Ig and non-Ig linked glycans to the trait value (see Supplementary Table 9), which was inferred based on information about protein-specific glycosylation reported previously in [9]. For more details about the procedure of Ig/non-Ig/mixed assignment see Supplementary Note 4.

**Figure 2.**
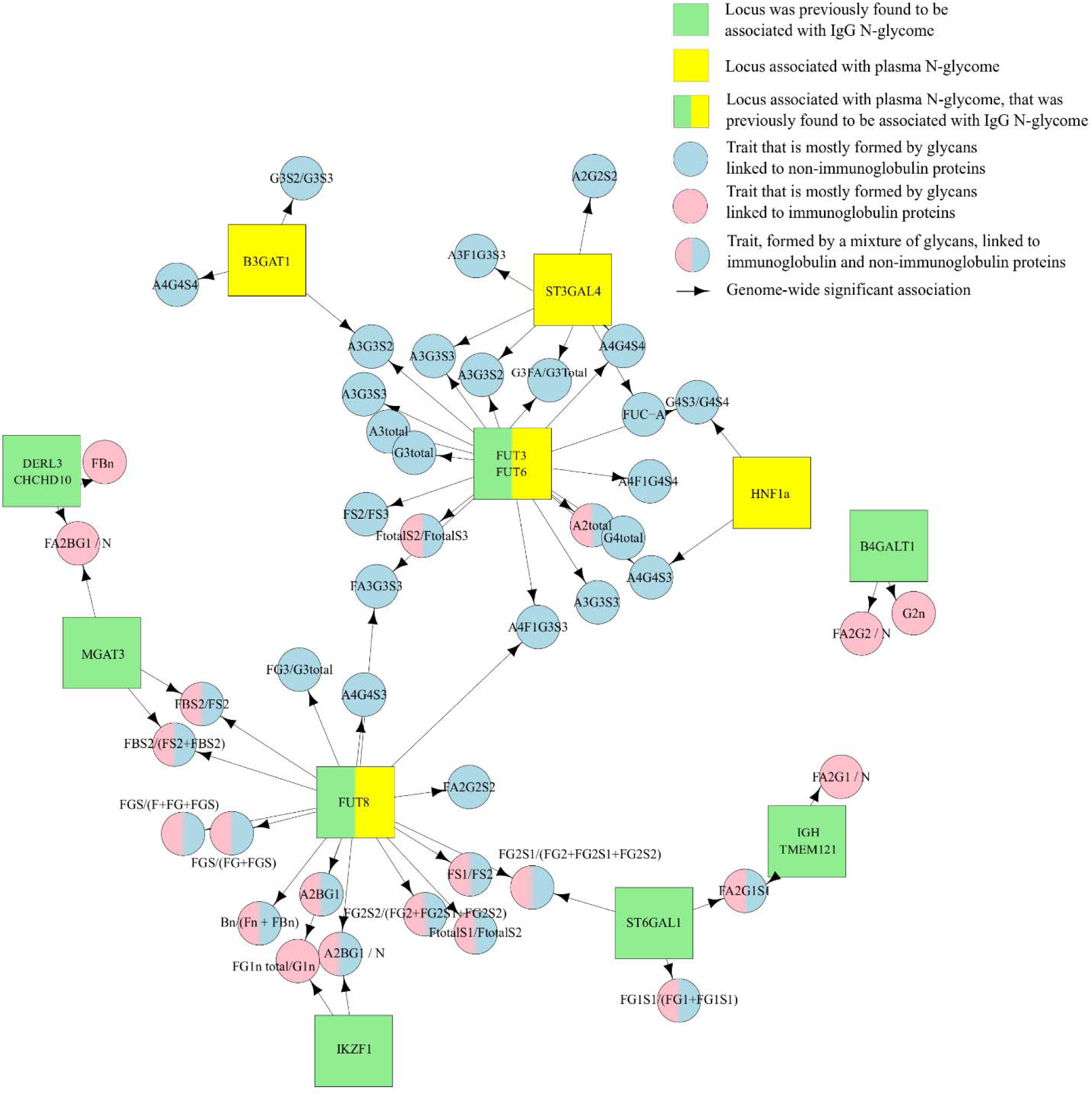
A network view of associations between loci and glycan traits. Square nodes represent genetic loci labeled with the names of candidate gene(s), circle nodes represent glycan traits. Green highlights candidate genes, located in genomic regions that were previously found to be associated with IgG N-glycome. Yellow highlights candidate genes, located in genomic regions associated with plasma N-glycome. Pink color highlights glycan traits mostly containing glycans that are linked to immunoglobulins. Blue color highlights traits that are mostly formed by glycans linked to other (not immunoglobulin) proteins. Blue/pink color highlights glycan traits, formed by a mixture of glycans that are linked to immunoglobulin and non-immunoglobulin proteins. Arrows represent genetic association (P-value ≤ 1.66 × 10^−9^) between gene and specific glycan.

The resulting network (Figure 2) shows that candidate genes and glycan traits cluster into two major subnetworks or hubs. The first subnetwork contained the six loci: *FUT8, DERL3/CHCHD10, IKZF1, TMEM121, ST6GAL1*, and *MGAT3*, with *FUT8* as a hub. These loci, as well as the locus containing *B4GATL1*, were associated with core-fucosylated biantennary glycans. It is known that the majority of plasma core-fucosylated biantennary glycans are linked to immunoglobulins [9]. Moreover, in previous studies these seven genes were found to be associated with N-glycosylation of IgG [13,48]. At the same time these genes were associated with non-immunoglobulins linked glycans. We can consider this cluster (seven genes out of eleven) as related to both IgG and non-IgG glycosylation. Taking into account that IgG is the most prevalent glycosylated plasma protein, it is not surprising that more than a half of replicated loci are actually associated with immunoglobulins glycosylation. However, previous GWAS on HPLC plasma N-glycome reported only one locus - *FUT8* - overlapping with IgG loci.

The second subnetwork in Figure 2 contained four loci (*ST3GAL4, HNF1a, FUT3/FUT6*, and *B3CAT1*, with *FUT3/FUT6* as a hub) associated with tri- and tetra-antennary glycans. It is known that these types of glycans are linked to plasma proteins other than IgG [9]. Thus we relate this cluster to non-IgG plasma protein N-glycosylation. Among these four loci we report *ST3CAL4* as the novel locus controlling the N-glycosylation of non-IgG plasma proteins. We attribute it to non-immunoglobulins plasma protein N-glycosylation owing to its association with tetra-antennary glycans.

## Discussion

We conducted the first genome-wide association study of total plasma N-glycome measured by UPLC technology. Our efforts brought the number of loci significantly associated with total plasma N-glycome from 6 [26,27] to 16, of which 12 were replicated in our work. This allowed us to next use a range of *in-silico* functional genomics analyses to identify candidate genes in the established loci and to obtain insight into biological mechanisms of plasma glycome regulation.

Compared to the HPLC glycan measurement technology used in previous GWAS of plasma N-glycome [26,27], UPLC technology provides better resolution and quantification of glycan structures, resulting in increased power of association testing: we have detected fourteen vs. six plasma N-glycome QTLs, despite the reduced sample size of our study (2.763 samples here vs. 3533 samples in [27]). It should be noted that we used new imputation panel (1000 Genomes instead of HapMap in the previous studies) that more than tripled the number of polymorphisms analyzed genome-wide (from 2.4M SNPs to 8M). That may have contributed to the higher power of our study as well. In addition to detecting novel loci, we were able to replicate five (*HNF1a, FUT6, FUT8, B3GAT1* and *MGAT5*) of six loci that were reported previously to be associated with human plasma N-glycome measured using the HPLC technology [26,27].

Among six plasma glycome loci that were identified as genome-wide significant previously [26,27], only one (regions of *FUT8*) had overlap with a locus identified as associated with IgG glycome composition [13]. A recent multivariate GWAS study of plasma IgG glycome composition [48] identified five new loci, including the region of *FUT3/FUT6*, thus bringing the overlap between plasma and IgG glycome loci to two. In our study, among 12 replicated loci, majority (eight) overlap with loci that were reported to be associated with IgG glycome composition [13,48] (see Figure 2). We therefore clearly establish a strong overlap between IgG and plasma glycome loci.

In a way, this overlap is to be expected. It is known that majority of serum (and therefore plasma) glycoproteins are either immunoglobulins produced by B-lymphocytes or glycoproteins secreted by the liver [53]. We thus expected overlap between IgG and total plasma glycome loci, and we expected that loci associated with the plasma N-glycome would be enriched by genes with tissue specific expression in liver and B-cells. Indeed, we find that plasma N-glycome loci are enriched for genes expressed in plasma cells, antibody producing cells and B-lymphocytes, and we also find overlap between plasma N-glycome loci and CD19+ eQTLs. However, we neither find enrichment of genes that are expressed in liver (Supplementary table 5c), nor overlap between plasma N-glycome loci and liver eQTLs. In the future, it will be important to achieve better resolution and separation of loci that are related to glycosylation of non-immunoglobulin glycoproteins. This could be achieved either technologically (e.g. performing analyses of IgG-free fractions of proteins), or this could be attempted via statistical modelling.

The genetic variation in the *FUT3/FUT6* locus is a major (in terms of proportion of variance explained and number of glycans affected) genetic factor for non-immunoglobulins glycosylation. According to current knowledge, these enzymes catalyze fucosylation of antennary GlcNAc32, resulting in glycan structures that are not found on IgG [9,54]. This is consistent with the spectrum of glycan traits associated with *FUT3/FUT6* locus in our work (Figure 2). However, this locus was recently found to be associated with IgG glycosylation [48]. The authors could not explain this finding because at that time IgG glycans were not known to contain antennary fucose. Two explanations could have been proposed for this surprising finding: either, enzymes encoded by *FUT3/FUT6* locus exhibit non-canonical activity of core fucosylation, or that some IgG glycans actually do contain antennary fucose. Recently, the latter was demonstrated in work by Russell et al. [14]. Our work does not show any evince for association between *FUT3/FUT6* locus and core fucosylation, hence providing an independent evidence that explanation of association between *FUT3/FUT6* and IgG glycosylation is rooted in presence of antennary fucose on some IgG glycans.

An interesting pattern starts emerging out of study of genetic control of plasma glycosylation. We now see a clear overlap in genetic control between plasma and IgG glycosylation, which calls for future studies that would help distinguish between global, cell-, tissue-, and protein-specific pathways of protein glycosylation. Many (eight out of twelve) replicated loci contained genes that encode enzymes directly involved in glycosylation (*FUT3/FUT6, FUT8, B3GAT1, ST6GAL1, B4GALT1, ST3GAL4, MGAT3*, and *MGAT5*). We, however, start now seeing loci and genes, which are likely to reflect other, more complex, aspects of plasma glycosylation process. These genes include *DERL3*, which potentially highlights the role of glycoprotein degradation pathway, and such transcription factors as *HNF1a* and *IKZF1*. Such regulatory genes, in our view, are plausible candidates that will help linking glycans and complex human disease. This view is supported by an example of mutations in *HNF1a*, that lead to maturity onset diabetes of the young (MODY), and to strong distortion of plasma glycosylation profile [24]. Further, bigger studies, using refined molecular and computational technologies, will allow expanding the list of genes involved in regulation of glycome composition, establish cell-, tissue-, and protein-specific glycosylation pathways and will substantiate and explain the relations between glycosylation and mechanisms of human health and disease. To facilitate further studies of glycosylation and of the role of glycome in human health and diseases we have made full results of our plasma N-glycome GWAS (almost one billion of trait-SNP associations) freely available to the scientific community via GWAS archive.

## Conclusion

Previous GWAS of HPLC measured plasma N-glycome [27] identified six genes controlling plasma N-glycosylation of which four implicated genes with obvious links to the glycosylation process. Here, using a smaller sample but more precise UPLC technology and new GWAS imputation panels, we confirmed the association of five known loci and identified and replicated additional seven loci. Our results demonstrate that genetic control of plasma proteins N-glycosylation is a complex process, which is under control of genes that belong to different pathways and are expressed in different tissues. Further studies with larger sample size should further decrypt the genetic architecture of the glycosylation process and explain the relations between glycosylation and mechanisms of human health and disease.

## Data availability

Summary statistics from our plasma N-glycome GWAS for 113 glycan traits are available for interactive exploration at the GWAS archive (http://gwasarchive.org). The data set was also deposited at Zenodo [55]. The data generated in the secondary analyses of this study are included with this article in the supplementary tables.

## Materials and Methods

### Study cohort description

This work is based on analysis of data from four cohorts - TwinsUK, PainOR, SOCCS and QMDiab. Sample demographics can be found in Supplementary Table 10.

### TwinsUK

The TwinsUK cohort [56] (also referred to as the UK Adult Twin Register) is an a nationwide registry of volunteer twins in the United Kingdom, with about 13,000 registered twins (83% female, equal number of monozygotic and dizygotic twins, predominantly middle-aged and older). The Department of TwinResearch and Genetic Epidemiology at King’s College London (KCL) hosts the registry. From this registry, a total of 2,763 subjects had N-linked total plasma glycan measurements which were included in the analysis.

### QMDiab

The Qatar Metabolomics Study on Diabetes (QMDiab) is a cross-sectional case–control study with 374 participants. QMDiab has been described previously and comprises male and female participants in near equal proportions, aged between 23 and 71 years, mainly of Arab, South Asian and Filipino descent [57,58]. The initial study was approved by the Institutional Review Boards of HMC and Weill Cornell Medicine—Qatar (WCM-Q) (research protocol #11131/11). Written informed consent was obtained from all participants. All study participants were enrolled between February 2012 and June 2012 at the Dermatology Department of Hamad Medical Corporation (HMC) in Doha, Qatar. Inclusion criteria were a primary form of type 2 diabetes (for cases) or an absence of type 2 diabetes (for controls). Sample collection was conducted in the afternoon, after the general operating hours of the morning clinic. Patient and control samples were collected in a random order as they became available and at the same location using identical protocols, instruments and study personnel. Samples from cases and controls were processed in the laboratory in parallel and in a blinded manner. Data from five participants were excluded from the analysis because of incomplete records, leaving 176 patients and 193 controls. Of the 193 control participants initially enrolled, 12 had HbA1c levels above 6.5% (48 mmol/mol) and were subsequently classified as cases, resulting in 188 cases and 181 controls.

### SOCCS

SOCCS study [59,60] comprised 2,057 (colorectal cancer) CRC cases (61% male; mean age at diagnosis 65.8±8.4 years) and 2,111 population controls (60% males; mean age 67.9±9.0 years) as ascertained in Scotland. Cases were taken from an independent, prospective, incident CRC case series and aged <80 years at diagnosis. Control subjects were population controls matched by age (±5 years), gender and area of residence within Scotland. All participants gave written informed consent and study approval was from the MultiCentre Research Ethics Committee for Scotland and Local Research Ethics committee. Sample collection is described in [59,60].

### PainOR

The PainOR [61] is the University of Parma cohort of patients of a retrospective multicenter study (ClinicalTrials.gov Identifier NCT02037789) part of the PainOMICS project funded by European Community in the Seventh Framework Programme (Project ID: 602736). The primary objective is to recognize genetic variants associated with chronic low back pain (CLBP); secondary objectives are to study glycomics and activomics profiles associated with CLBP. Glycomic and Activomic approaches aim to reveal alterations in proteome complexity that arise from post-translational modification that varies in response to changes in the physiological environment, a particularly important avenue to explore in chronic inflammatory diseases. The study was firstly approved by the Institutional Review Boards of IRCCS Foundation San Matteo Hospital Pavia and then by the Institutional Review boards of all clinical centers that enrolled patients. Copies of approvals were provided to the European Commission before starting the study. Written informed consent was obtained from all participants. In the period between September 2014 and February 2016, one thousand of patients (including 38.1% male and 61.9% female, averaging 65±14.5 years) were enrolled at the Anesthesia, Intensive Care and Pain Therapy Department of University Parma Hospital. Inclusion criteria were adult Caucasian patients who were suffering of low back pain (pain between the costal margins and gluteal fold, with or without symptoms into one or both legs) more than 3 months who were admitted at Pain Department of University Parma Hospital. We exclude patients with recent history of spinal fractures or low back pain due to cancer or infection. Sample collection was performed in all patients enrolled, according to the Standard Operating Procedures published in PlosOne in 2017 [62]. Samples were processed in PainOmics laboratory in a blinded manner in University of Parma.

### Genotyping

For full details of the genotyping and imputation see Supplementary Table 11.

### TwinsUK

Genotyping was carried out using combination Illumina SNP arrays: HumanHap300, HumanHap610Q, 1M - Duo and 1.2MDuo 1M. Standard quality control of genotyped data was applied, with SNPs filtered by sample call rate > 98%, MAF > 1%, SNP call rate: >97% (for SNP with MAF≥5%) or >99% (for SNPs with 1% ≤ MAF <5%), HWE P-value ≤ 1 × 10^−6^. In total 275,139 SNPs passed criteria. Imputation was done using IMPUTE2 software with 1000G phase 1 version 3 and mapped to the GRCh37 human genome build. Imputed SNPs were filtered by imputation quality (SNPTEST proper-info) > 0.7., MAF >= 1%; MAC >= 10; leading to 8,557,543 SNPs passed to the GWAS analysis.

### QMDiab

Genotyping was carried out using Illumina Omni array 2.5 (version 8). Standard quality control of genotyped data was applied, with SNPs filtered by sample call rate > 98%, MAF > 1%, SNP call rate: > 98%, HWE P-value ≤ 1 × 10^−6^. In total 1,223,299 SNPs passed criteria. Imputation was done using SHAPEIT software with 1000G phase 3 version 5 and mapped to the GRCh37 human genome build. Imputed SNPs were filtered by imputation quality > 0.7, leading to 20,483,276 SNPs passed to the GWAS analysis.

### SOCCS

Details of the genotyping procedure can be found here [63]. Genotyping was carried out using Illumina SNP arrays: HumanHap300 and HumanHap240S. Standard quality control of genotyped data was applied, with SNPs filtered by sample call rate > 95%, MAF > 1%, SNP call rate: >95%, HWE P-value ≤ 1 × 10^−6^. In total 514,177 SNPs passed criteria. Imputation was done using SHAPEIT and IMPUTE2 software with 1000 Genomes, phase 1 (Integrated haplotypes, released June 2014) and mapped to the GRCh37 human genome build. Imputed SNPs were not filtered, leading to 37,780,221 SNPs passed to the GWAS analysis.

### PainOR

Genotyping was carried out using Illumina HumanCore BeadChip. Standard quality control of genotyped data was applied with SNPs filtered by sample call rate >98%, MAF >0.625%, SNP call rate: > 97%, HWE P-value ≤ 1 × 10^−6^. In total 253,149 SNPs passed criteria. Imputation was done using Eagle software with HRC r1.1 2016 reference and mapped to the GRCh37 human genome build. Imputed SNPs were not filtered, leading to 39,127,685 SNPs passed to the GWAS analysis.

### Phenotyping

#### Plasma N-glycome quantification

Plasma N-glycome quantification of samples from TwinsUK, PainOR and QMDiab were performed at Genos by applying the following protocol. Plasma N-glycans were enzymatically released from proteins by PNGase F, fluorescently labelled with 2-aminobenzamide and cleaned-up from the excess of reagents by hydrophilic interaction liquid chromatography solid phase extraction (HILIC-SPE), as previously described. [64]. Fluorescently labelled and purified N-glycans were separated by HILIC on a Waters BEH Glycan chromatography column, 150 × 2.1 mm, 1.7 μm BEH particles, installed on an Acquity ultra-performance liquid chromatography (UPLC) instrument (Waters, Milford, MA, USA) consisting of a quaternary solvent manager, sample manager and a fluorescence detector set with excitation and emission wavelengths of 250 nm and 428 nm, respectively. Following chromatography conditions previously described in details [64], total plasma N-glycans were separated into 39 peaks for QMDiab, TwinsUK and PainOR cohorts. The amount of N-glycans in each chromatographic peak was expressed as a percentage of total integrated area. Glycan peaks (GPs) - quantitative measurements of glycan levels - were defined by automatic integration of intensity peaks on chromatogram. Number of defined glycan peaks varied among studies from 36 to 42 GPs.

Plasma N-glycome quantification for SOCCS samples were done at NIBRT by applying the same protocol as for TwinsUK, PainOR and QMDiab, with the only difference in the excitation wavelength (330 nm instead of 250 nm).

#### Harmonization of glycan peaks

The order of the glycan peaks on a UPLC chromatogram was similar among the studies. However, depending on the cohort some peaks located near one another might have been indistinguishable. The number of defined glycan peaks (GPs) varied among studies from 36 to 42. To conduct GWAS on TwinsUK following by replication in other cohorts, we harmonized the set of peaks (or GPs). According to the major glycostructures within the GPs we manually created the table of correspondence between different GPs (or sets of GPs) across all cohorts, where plasma glycome was measured using UPLC technology. Then, based on this table of correspondence, we defined the list of 36 harmonized GPs (Supplementary Table 12) and the harmonization scheme for each cohort. We validated the harmonization protocol by comparing with manual re-integration of the peaks on chromatogram level using 35 randomly chosen samples from 3 cohorts: TwinsUK, PainOR and QMDiad. We show the full concordance between two approaches (Pearson correlation coefficient R>0.999, see Supplementary Table 12 for the details). We applied this harmonization procedure for the four cohorts: TwinsUK, QMDiab, CRC and PainOR, leading to the set of 36 glycan traits in each cohort.

#### Normalization and batch-correction of GPs

Normalization and batch-correction was performed on harmonized UPLC glycan data for four cohorts: TwinsUK, PainOR, SOCCS and QMDiab. We used total area normalization (the area of each GP was divided by the total area of the corresponding chromatogram). Normalized glycan measurements were log10-transformed due to right skewness of their distributions and the multiplicative nature of batch effects. Prior to batch correction, samples with outlying measurements were removed. Outlier was defined as a sample that had at least one GP that is out of 3 standard deviation from the mean value of GP. Batch correction was performed on log10-transformed measurements using the ComBat method, where the technical source of variation (batch and plate number) was modelled as a batch covariate. Again, samples with outlying measurements were removed.

From the 36 directly measured glycan traits, 77 derived traits were calculated (see Supplementary Table 9). These derived traits average glycosylation features such as branching, galactosylation and sialylation across different individual glycan structures and, consequently, they may be more closely related to individual enzymatic activity and underlying genetic polymorphism. As derived traits represent sums of directly measured glycans, they were calculated using normalized and batch-corrected glycan measurements after transformation to the proportions (exponential transformation of batch-corrected measurements). The distribution of 113 glycan traits can be found in Supplementary Figure 2.

Prior to GWAS, the traits were adjusted for age and sex by linear regression. The residuals were rank transformed to normal distribution (rntransform function in GenABEL [65,66] R package).

#### Genome-wide association analysis

Discovery GWAS was performed using TwinsUK cohort (N = 2,763) for 113 GP traits. GEMMA [67] was used to estimate the kinship matrix and to run linear mixed model regression on SNP dosages assuming additive genetic effects. Obtained summary statistics were corrected for genomic control inflation factor λ_GC_ to account for any residual population stratification. An association was considered statistically significant at the genome-wide level if the P-value for an individual SNP was less than 5 × 10^−8^ / (29+1) = 1.66 × 10^−9^, where 29 is an effective number of tests (traits) that was estimated as the number of principal components that jointly explained 99% of the total plasma glycome variance in the TwinsUK sample.

#### Locus definition

In short, we considered SNPs located in the same locus if they were located within 500 Kb from the leading SNP (the SNP with lowest P-value). Only the SNPs and the traits with lowest P-values are reported (leading SNP-trait pairs). The detailed procedure of locus definition is described in Supplementary Note 1.

#### Replication

We have used TwinsUK cohort for the replication of six previously described loci [27] affecting plasma N-glycome. From each of six loci we have chosen leading SNP with the strongest association as reported by authors [27]. Since there is no direct trait-to-trait correspondence between glycan traits measured by HPLC and UPLC technologies we tested the association of the leading SNPs with all 113 PGPs in TwinsUK cohort. We considered locus as replicated if its leading SNP showed association with at least one of 113 PGPs with replication threshold of P-value≤0.05/(6*30) = 2.78 × 10^−4^, where six is number of loci and 30 is a number of principal components that jointly explained 99% of the total plasma N-glycome variance.

For the replication of novel associations, we used data from 3 cohorts: PainOR (N = 294), QMDiab (N = 327) and SOCCS (N = 472) with total replication sample size of N = 1,048 samples that have plasma UPLC N-glycome and genotype data (for details of genotyping, imputation and association analysis, see Supplementary Table 11). We used only the leading SNPs and traits for the replication that were identified in the discovery step. For these SNPs we conducted a fixed-effect meta-analysis using METAL software [68] combining association results from three cohorts. The replication threshold was set as P-value≤0.05/10=0.005, where 10 is the number of replicated loci. Moreover, we checked whether the sign of estimated effect was concordant between discovery and replication studies.

### Functional annotation *in-silico*

#### Variant effect prediction (VEP)

For annotation with the variant effect predictor (VEP, [32]), for each of the 12 replicated loci we have selected the set of SNPs that had strong associations, defined as those located within +/− 250kbp window from the strongest association, and having P-value≤T, where log10(T)=log10(P_min_)+1, where P_min_ is the P-value of the strongest association in the locus.

#### Gene-set and tissue/cell enrichment analysis

To prioritize genes in associated regions, gene set enrichment and tissue/cell type enrichment analyses were carried out using DEPICT software v. 1 rel. 194 [35]. For the analysis we have chosen independent variants (see “Locus definition”) with P-value ≤5 × 10^−8^/30 (14 SNPs) and P-value <1×10^−5^/30 (93 SNPs). We used 1000G data set for calculation of LD [69]s.

#### Pleiotropy with complex traits

We have investigated the overlap between associations obtained here and elsewhere, using PhenoScanner v1.1 database [36]. For twelve replicated SNPs (Table 1, Table 2) we looked up traits that have demonstrated genome-wide significant (p < 5 × 10−8) association at the same or at strongly (r2 < 0.7) linked SNPs.

#### Pleiotropy with eQTLs

To identify genes whose expression levels could potentially mediate the association between SNPs and plasma glycan traits we performed a summary-data based Mendelian randomization (SMR) analysis followed by heterogeneity in dependent instruments (HEIDI) method [44]. In short, SMR test aims at testing the association between gene expression (in a particular tissue) and a trait using the top associated expression quantitative trait loci (eQTL) as a genetic instrument. Significant SMR test indicates evidence of causality or pleiotropy but also the possibility that SNPs controlling gene expression are in linkage disequilibrium with those associated with the traits. These two situations can be disentangled using the HEIDI (HEterogeneity In Dependent Instrument) test.

The SMR/HEIDI analysis was carried out for leading SNPs that were replicated and were genome-wide significant (P-value≤1.7 × 10^−9^) on discovery stage (11 loci in total, see Table 1). We checked for overlap between these loci and eQTLs in blood [45], 44 tissues provided by the GTEx database [46] and in 9 cell lines from CEDAR dataset [47], including six circulating immune cell types (CD4+ T-lymphocytes, CD8+ T lymphocytes, CD19+ B lymphocytes, CD14+ monocytes, CD15+ granulocytes, platelets. Technical details of the procedure may be found in Supplementary Note 2. Following Bonferroni procedure, the results of the SMR test were considered statistically significant if P-value_SMR_ < 2.445 × 10^−6^ (0.05/20448, where 20448 is a total number of probes used in analysis for all three data sets). Inferences whether functional variant may be shared between plasma glycan trait and expression were made based on HEIDI test: p > 0.05 (likely shared), 0.05 > p > 0.001 (possibly shared), p < 0.001 (sharing is unlikely).

## Acknowledgments

This work was supported by the European Community’s Seventh Framework Programme funded project PainOmics (Grant agreement # 602736) and by the European Structural and Investments funding for the “Croatian National Centre of Research Excellence in Personalized Healthcare” (contract #KK. 01.1.1.01.0010).The work of SSh was supported by the Russian Ministry of Science and Education under the 5–100 Excellence Programme.

The work of YT and YA was supported by the Federal Agency of Scientific Organizations via the Institute of Cytology and Genetics (project #0324-2018-0017).

Karsten Suhre and Gaurav Thareja are supported by ‘Biomedical Research Program’ funds at Weill Cornell Medicine - Qatar, a program funded by the Qatar Foundation. We thank all staff at Weill Cornell Medicine - Qatar and Hamad Medical Corporation, and especially all study participants who made the QMDiab study possible.

The SOCCS study was supported by grants from Cancer Research UK (C348/A3758, C348/A8896, C348/ A18927); Scottish Government Chief Scientist Office (K/OPR/2/2/D333, CZB/4/94); Medical Research Council (G0000657-53203, MR/K018647/1); Centre Grant from CORE as part of the Digestive Cancer Campaign (http://www.corecharity.org.uk).

TwinsUK is funded by the Wellcome Trust, Medical Research Council, European Union, the National Institute for Health Research (NIHR)-funded BioResource, Clinical Research Facility and Biomedical Research Centre based at Guy’s and St Thomas’ NHS Foundation Trust in partnership with King’s College London.

## Author Contributions

SSh and YT contributed to the design of the study, carried out statistical analysis, produced the figures; SSh, YT, LK, KS, YA produced wrote the manuscript; LK, FV, SSh, JK contributed to data harmonization and quality control; MS, MV, FV, TP, JerS, ITA, JK, JelS, MPB, GL contributed to plasma N-glycome measurements; MM and TS analyzed TwinsUK dataset and contributed to interpretation of the results; LK, AM, HC, MD, SF analyzed SOCCS dataset and contributed to interpretation of the results; MA, FW and CD designed PainOR study and contributed to interpretation of the results; KS and GT analyzed QMDiab dataset and contributed to interpretation of the results; EL, JD and MG designed CEDAR study and contributed to interpretation of the results; YA and GL conceived and oversaw the study, contributed to the design and interpretation of the results; all co-authors contributed to the final manuscript revision.

## Supplementary Information

Supplementary Figure 1 – Quantile-quantile (QQ) plots of association analysis of 113 PGPs

Supplementary Figure 2 – Distribution of 113 PGPs measured for 2763 TwinsUK samples after correction for sex and age

Supplementary Note 1 – Locus definition

Supplementary Note 2 – Testing for pleiotropic effects using SMR/HEIDI approach

Supplementary Note 3 – Correlated Expression & Disease Association Research (CEDAR)

Supplementary Note 4 – Classification of glycans into Ig-related and non-Ig

Supplementary Table 1 – Replication of previously published associations with plasma N– glycome.

Supplementary Table 2 – Genomic control inflation factor lambda for 113 traits (discovery GWAS)

Supplementary Table 3 – Discovery and replication of fourteen loci associated with plasma N– glycome (P-value < 1.66e-9)

Supplementary Tables 4 – Results of VEP analysis for the most associated SNPs for twelve replicated loci

Supplementary Table 5 – Results of DEPICT analysis for significant loci (1.7e-9)

Supplementary Table 6 – Results of DEPICT analysis for suggestively significant loci (3.3e-7)

Supplementary Table 7 – Results of PhenoScaner analysis

Supplementary Table 8 – Results of SMR-HEIDI analysis

Supplementary Table 9 – The description of 36 quantitative plasma N-glycosylation traits measured by UPLC and 77 derived traits

Supplementary Table 10 – Sample demographics

Supplementary Table 11 – Details of the genotyping, imputation and association analysis for studied cohorts

Supplementary Table 12 – Correspondence between glycan peaks (GP), obtained in the following studies: TwinsUK, FinRisk, Dundee, QMDiab, PainOR, SABRE, SOCCS

Supplementary Table 13 – Comparison of area summation approach with manual integration approach

## References

1. Varki A. Biological roles of oligosaccharides: all of the theories are correct // Glycobiology. 1993. Vol. 3, № 2. P. 97–130.

2. Ohtsubo K., Marth J.D. Glycosylation in Cellular Mechanisms of Health and Disease // Cell. 2006. Vol. 126, № 5. P. 855–867.

3. Skropeta D. The effect of individual N-glycans on enzyme activity // Bioorg. Med. Chem. 2009. Vol. 17, № 7. P. 2645–2653.

4. Takeuchi H. et al. O-Glycosylation modulates the stability of epidermal growth factor-like repeats and thereby regulates Notch trafficking. // J. Biol. Chem. American Society for Biochemistry and Molecular Biology, 2017. Vol. 292, № 38. P. 15964–15973.

5. Lauc G. et al. Mechanisms of disease: The human N-glycome. // Biochim. Biophys. Acta. Elsevier, 2015. Vol. 1860, № 8. P. 1574–1582.

6. Poole J. et al. Glycointeractions in bacterial pathogenesis // Nat. Rev. Microbiol. Nature Publishing Group, 2018. P. 1.

7. Khoury G.A., Baliban R.C., Floudas C.A. Proteome-wide post-translational modification statistics: frequency analysis and curation of the swiss-prot database. // Sci. Rep. Nature Publishing Group, 2011. Vol. 1.

8. Craveur P., Rebehmed J., de Brevern A.G. PTM-SD: a database of structurally resolved and annotated posttranslational modifications in proteins. // Database (Oxford). Oxford University Press, 2014. Vol. 2014.

9. Clerc F. et al. Human plasma protein N-glycosylation // Glycoconj. J. Springer, 2016. Vol. 33, № 3. P. 309–343.

10. Freidin M.B. et al. The Association Between Low Back Pain and Composition of IgG Glycome. // Sci. Rep. Nature Publishing Group, 2016. Vol. 6. P. 26815.

11. Gudelj I. et al. Low galactosylation of IgG associates with higher risk for future diagnosis of rheumatoid arthritis during 10 years of follow-up // Biochim. Biophys. Acta—Mol. Basis Dis. 2018. Vol. 1864, № 6. P. 2034–2039.

12. Connelly M.A. et al. Inflammatory glycoproteins in cardiometabolic disorders, autoimmune diseases and cancer // Clinica Chimica Acta. 2016. Vol. 459. P. 177–186.

13. Lauc G. et al. Loci associated with N-glycosylation of human immunoglobulin G show pleiotropy with autoimmune diseases and haematological cancers. // PLoS Genet. / ed. Gibson G. 2013. Vol. 9, № 1. P. e1003225.

14. Russell A.C. et al. The N-glycosylation of immunoglobulin G as a novel biomarker of Parkinson’s disease. // Glycobiology. 2017. Vol. 27, № 5. P. 501–510.

15. Lemmers R.F.H. et al. IgG glycan patterns are associated with type 2 diabetes in independent European populations // Biochim. Biophys. Acta—Gen. Subj. 2017. Vol. 1861, 9. P. 2240–2249.

16. Freeze H.H. Genetic defects in the human glycome // Nat. Rev. Genet. Nature Publishing Group, 2006. Vol. 7, № 7. P. 537–551.

17. Trbojevic Akmacic I. et al. Inflammatory bowel disease associates with proinflammatory potential of the immunoglobulin G glycome. // Inflamm. Bowel Dis. Wolters Kluwer Health, 2015. Vol. 21, № 6. P. 1237–1247.

18. Fuster M.M., Esko J.D. The sweet and sour of cancer: glycans as novel therapeutic targets // Nat. Rev. Cancer. Nature Publishing Group, 2005. Vol. 5, № 7. P. 526–542.

19. Dube D.H., Bertozzi C.R. Glycans in cancer and inflammation — potential for therapeutics and diagnostics // Nat. Rev. Drug Discov. Nature Publishing Group, 2005. Vol. 4, № 6. P. 477–488.

20. Pagan J.D., Kitaoka M., Anthony R.M. Engineered Sialylation of Pathogenic Antibodies In Vivo Attenuates Autoimmune Disease. // Cell. 2018. Vol. 172, № 3. P. 564–577.e13.

21. Adamczyk B., Tharmalingam T., Rudd P.M. Glycans as cancer biomarkers. // Biochim. Biophys. Acta. 2012. Vol. 1820, № 9. P. 1347–1353.

22. Maverakis E. et al. Glycans in the immune system and The Altered Glycan Theory of Autoimmunity: a critical review. // J. Autoimmun. 2015. Vol. 57. P. 1–13.

23. RodrÍguez E., Schetters S.T.T., van Kooyk Y. The tumour glyco-code as a novel immune checkpoint for immunotherapy. // Nat. Rev. Immunol. 2018. Vol. 18, № 3. P. 204–211.

24. Thanabalasingham G. et al. Mutations in HNF1A result in marked alterations of plasma glycan profile. // Diabetes. 2013. Vol. 62, № 4. P. 1329–1337.

25. Taniguchi N., Kizuka Y. Glycans and cancer: role of N-glycans in cancer biomarker, progression and metastasis, and therapeutics. // Adv. Cancer Res. 2015. Vol. 126. P. 11–51.

26. Lauc G. et al. Genomics meets glycomics-the first GWAS study of human N-Glycome identifies HNF1α as a master regulator of plasma protein fucosylation. // PLoS Genet. 2010. Vol. 6, № 12. P. e1001256.

27. Huffman J.E. et al. Polymorphisms in B3GAT1, SLC9A9 and MGAT5 are associated with variation within the human plasma N-glycome of 3533 European adults. // Hum. Mol. Genet. Oxford University Press, 2011. Vol. 20, № 24. P. 5000–5011.

28. Huffman J.E. et al. Comparative performance of four methods for high-throughput glycosylation analysis of immunoglobulin G in genetic and epidemiological research. // Mol. Cell. Proteomics. 2014. Vol. 13, № 6. P. 1598–1610.

29. Knežević A. et al. High throughput plasma N-glycome profiling using multiplexed labelling and UPLC with fluorescence detection // Analyst. The Royal Society of Chemistry, 2011. Vol. 136, № 22. P. 4670.

30. 1000 Genomes Project Consortium {fname} et al. A map of human genome variation from population-scale sequencing. // Nature. 2010. Vol. 467, № 7319. P. 1061–1073.

31. Consortium the H.R. et al. A reference panel of 64,976 haplotypes for genotype imputation // Nat. Genet. Nature Publishing Group, 2016. Vol. 48, № 10. P. 1279–1283.

32. McLaren W. et al. The Ensembl Variant Effect Predictor. // Genome Biol. 2016. Vol. 17, № 1. P. 122.

33. Adzhubei I.A. et al. A method and server for predicting damaging missense mutations. // Nat. Methods. 2010. Vol. 7, № 4. P. 248–249.

34. Kumar P., Henikoff S., Ng P.C. Predicting the effects of coding non-synonymous variants on protein function using the SIFT algorithm // Nat. Protoc. 2009. Vol. 4, № 7. P. 1073–1081.

35. Pers T.H. et al. Biological interpretation of genome-wide association studies using predicted gene functions // Nat. Commun. Nature Publishing Group, 2015. Vol. 6, № 1. P. 5890.

36. Staley J.R. et al. PhenoScanner: a database of human genotype-phenotype associations // Bioinformatics. 2016. Vol. 32, № 20. P. 3207–3209.

37. Teslovich T.M. et al. Biological, clinical and population relevance of 95 loci for blood lipids // Nature. 2010. Vol. 466, № 7307. P. 707–713.

38. Willer C.J. et al. Discovery and refinement of loci associated with lipid levels // Nat. Genet. 2013. Vol. 45, № 11. P. 1274–1283.

39. Shah T. et al. Gene-Centric Analysis Identifies Variants Associated With Interleukin-6 Levels and Shared Pathways With Other Inflammation Markers // Circ. Cardiovasc. Genet. 2013. Vol. 6, № 2. P. 163–170.

40. Ridker P.M. et al. Loci related to metabolic-syndrome pathways including LEPR, HNF1A, IL6R, and GCKR associate with plasma C-reactive protein: the Women’s Genome Health Study. // Am. J. Hum. Genet. 2008. Vol. 82, № 5. P. 1185–1192.

41. Chambers J.C. et al. Genome-wide association study identifies loci influencing concentrations of liver enzymes in plasma // Nat. Genet. Nature Publishing Group, 2011. Vol. 43, № 11. P. 1131–1138.

42. Wood A.R. et al. Defining the role of common variation in the genomic and biological architecture of adult human height // Nat. Genet. Nature Publishing Group, a division of Macmillan Publishers Limited. All Rights Reserved., 2014. Vol. 46, № 11. P. 1173–1186.

43. Perry J.R.B. et al. Parent-of-origin-specific allelic associations among 106 genomic loci for age at menarche // Nature. 2014. Vol. 514, № 7520. P. 92–97.

44. Zhu Z. et al. Integration of summary data from GWAS and eQTL studies predicts complex trait gene targets // Nat. Genet. Nature Publishing Group, 2016. Vol. 48, № 5. P. 481–487.

45. Westra H.-J. et al. Systematic identification of trans eQTLs as putative drivers of known disease associations. // Nat. Genet. Nature Publishing Group, a division of Macmillan Publishers Limited. All Rights Reserved., 2013. Vol. 45, № 10. P. 1238–1243.

46. GTEx Consortium et al. Genetic effects on gene expression across human tissues. // Nature. 2017. Vol. 550, № 7675. P. 204–213.

47. Momozawa Y. et al. IBD risk loci are enriched in multigenic regulatory modules encompassing putative causative genes. // Nat. Commun. Nature Publishing Group, 2018. Vol. 9, № 1. p. 2427.

48. Shen X. et al. Multivariate discovery and replication of five novel loci associated with Immunoglobulin G N-glycosylation // Nat. Commun. Nature Publishing Group, 2017. Vol. 8, № 1. P. 447.

49. Pottier N. et al. Expression of SMARCB1 modulates steroid sensitivity in human lymphoblastoid cells: identification of a promoter snp that alters PARP1 binding and SMARCB1 expression // Hum. Mol. Genet. 2007. Vol. 16, № 19. P. 2261–2271.

50. Oda Y. et al. Derlin-2 and Derlin-3 are regulated by the mammalian unfolded protein response and are required for ER-associated degradation // J. Cell Biol. 2006. Vol. 172, № 3. P. 383–393.

51. Ardlie K.G. et al. The Genotype-Tissue Expression (GTEx) pilot analysis: Multitissue gene regulation in humans // Science (80-.). American Association for the Advancement of Science, 2015. Vol. 348, № 6235. P. 648–660.

52. Slack J.M.W. Molecular Biology of the Cell // Principles of Tissue Engineering. Garland Science, 2014. P. 127–145.

53. Bekesova S. et al. N-glycans in liver-secreted and immunoglogulin-derived protein fractions. // J. Proteomics. NIH Public Access, 2012. Vol. 75, № 7. P. 2216–2224.

54. Ma B., Simala-Grant J.L., Taylor D.E. Fucosylation in prokaryotes and eukaryotes // Glycobiology. 2006. Vol. 16, № 12. P. 158R–184R.

55. Sharapov S. et al. Genome-wide association summary statistics for human blood plasma glycome. 2018.

56. Moayyeri A. et al. The UK Adult Twin Registry (TwinsUK Resource) // Twin Res. Hum. Genet. Cambridge University Press, 2013. Vol. 16, № 01. P. 144–149.

57. Mook-Kanamori D.O. et al. 1,5-Anhydroglucitol in saliva is a noninvasive marker of short-term glycemic control. // J. Clin. Endocrinol. Metab. 2014. Vol. 99, № 3. P. E479–83.

58. Suhre K. et al. Connecting genetic risk to disease end points through the human blood plasma proteome // Nat. Commun. Nature Publishing Group, 2017. Vol. 8. P. 14357.

59. Vuckovic F. et al. IgG glycome in colorectal cancer // Clin. Cancer Res. American Association for Cancer Research, 2016. Vol. 22, № 12. P. 3078–3086.

60. Theodoratou E. et al. Glycosylation of plasma IgG in colorectal cancer prognosis // Sci. Rep. Nature Publishing Group, 2016. Vol. 6, № 1. P. 28098.

61. Allegri M. et al. ‘Omics’ biomarkers associated with chronic low back pain: protocol of a retrospective longitudinal study // BMJ Open. British Medical Journal Publishing Group, 2016. Vol. 6, № 10. P. e012070.

62. Dagostino C. et al. Validation of standard operating procedures in a multicenter retrospective study to identify-omics biomarkers for chronic low back pain // PLoS One / ed. Samartzis D. 2017. Vol. 12, № 5. P. e0176372.

63. Tenesa A. et al. Genome-wide association scan identifies a colorectal cancer susceptibility locus on 11q23 and replicates risk loci at 8q24 and 18q21 // Nat. Genet. Nature Publishing Group, 2008. Vol. 40, № 5. P. 631–637.

64. Akmačić I.T. et al. High-throughput glycomics: optimization of sample preparation. // Biochem. Biokhimii□a□. 2015. Vol. 80, № 7. P. 934–942.

65. Aulchenko Y.S. et al. GenABEL: an R library for genome-wide association analysis. // Bioinformatics. 2007. Vol. 23, № 10. P. 1294–1296.

66. Karssen L.C., van Duijn C.M., Aulchenko Y.S. The GenABEL Project for statistical genomics // F1000Research. 2016. Vol. 5. P. 914.

67. Zhou X., Stephens M. Efficient multivariate linear mixed model algorithms for genome-wide association studies // Nat. Methods. 2014. Vol. 11, № 4. P. 407–409.

68. Willer C.J., Li Y., Abecasis G.R. METAL: fast and efficient meta-analysis of genomewide association scans. // Bioinformatics. 2010. Vol. 26, № 17. P. 2190–2191.

69. Auton A. et al. A global reference for human genetic variation // Nature. 2015. Vol. 526, № 7571.

